# Scalable probabilistic matrix factorization for single-cell RNA-seq analysis

**DOI:** 10.1101/496810

**Authors:** Pedro F. Ferreira, Alexandra M. Carvalho, Susana Vinga

## Abstract

**Motivation:** The gene expression profile of a cell dictates its function in molecular processes, and can be used to probe its health status. This represents a step forward in the deep characterization of diseases such as cancer and may lead to breakthroughs in their treatment. The technology used to measure the gene expression of isolated cells, single-cell RNA-seq (scRNA-seq), has emerged in the last decade as a key enabler of this progress. However, the use of existing methods for dimensionality reduction, clustering and differential expression is limited by the specificities of the data obtained from scRNA-seq experiments, where technical factors may confound analyses of the true biological signal and contribute to spurious results. To overcome this issue, a possible approach is designing probabilistic generative models of the data with hidden variables encoding different underlying processes.

**Results:** We propose two novel probabilistic models for scRNA-seq data: modified probabilistic count matrix factorization (m-pCMF) and Bayesian zero-inflated negative binomial factorization (ZINBayes). These build upon previous models in the literature while leveraging scalable Bayesian inference via variational methods. We show that the proposed methods are competitive with the state-of-the-art models for robust dimensionality reduction in modern data sets, and improve upon the current best Bayesian model for small numbers of cells. The results show that building probabilistic models of latent variables which encode domain knowledge and using variational inference constitute a promising approach to analyse scRNA-seq data in a scalable way.

**Availability:** m-pCMF and ZINBayes are publicly available as Python packages at https://github.com/pedrofale/, along with the code to reproduce all the results.

## 1 Introduction

Single-cell RNA-sequencing (scRNA-seq) has emerged in the last decade as a key technology in using gene expression to study cell heterogeneity (Kolodziejczyk *et al.*, 2015). With the data obtained from these experiments, researchers can, for example, apply clustering algorithms to identify cell types and find genes which are differentially expressed between two conditions.

In scRNA-seq, the expression of a gene in a cell is measured by counting the number of mRNA molecules. Formally, let *N* be the number of cells in a data set and P the number of genes. Then, the expression matrix **X** is of size *N* × *P*, where each observation *x*_*np*_ contains the counts of mRNA molecules per gene *p* in cell *n*. The observations are assumed to be independent across both cells and genes.

Due to the large number of genes measured in a scRNA-seq experiment (never less than a few hundreds and occasionally up to tens of thousands, depending on the data acquisition protocol), the most common initial step in scRNA-seq data analysis is dimensionality reduction, i.e., reducing the observations from the original space ℕ^*P*^ to a lower-dimensional one, thus obtaining a reduced data matrix **Z** of size *N* × *K*, with *K* < *P*. This makes clustering analyses more tractable and, if *K* = 2 or *K* = 3, it allows for easy data visualization.

However, because mRNA molecules are counted within single cells, the total number of transcripts available may be very small and so they may not be detected at all. These undetected transcripts - often called “dropouts” - result in more zero counts than expected in the data matrix (Dal Molin *et al.*, 2017). In practice, this means there are two types of zeroes in **X**: some due to a true lack of expression and others due to dropouts. Although there are other confounding factors which deserve attention - capture efficiency and sequencing depth, batch effects and transcriptional noise –, this ambiguity makes dropouts the strongest source of noise in the data (Vallejos *et al.*, 2017).

As the mentioned confounders hide the true biological variability, using traditional methods for dimensionality reduction, clustering or differential expression is not reliable. To overcome this, several probabilistic models for scRNA-seq data have been proposed, along with corresponding inference mechanisms: ZIFA (Pierson and Yau, 2015), pCMF (Durif *et al.*, 2018), ZINB-WaVE (Risso *et al.*, 2018) and scVI (Lopez *et al.*, 2018).

The present paper considers the problem of performing scalable inference on useful scRNA-seq models to perform accurate downstreamanalyses, mainly clustering of cell types in big data sets. We propose two novel probabilistic models for scRNA-seq data: modified probabilistic count matrix factorization (m-pCMF) and Bayesian zero-inflated negative binomial factorization (ZINBayes). These build upon previous models in the literature while leveraging scalable Bayesian inference via variational methods. We compare their performance with existing models on a set of benchmarking tasks.

The remaining of this document is organized as follows. Section 2 introduces the proposed models and Section 3 assesses their performance on real public data sets. In Section 4 conclusions are drawn.

## 2 Methods

### 2.1 m-pCMF

Our first proposal is a modified version of pCMF (Durif *et al.*, 2018) in the following way:

- Introduce cell-specific scaling factors, similarly to ZINB-WaVE and scVI. These are meant to account for capture efficiency and sequencing depth variations.
- Introduce batch index annotations, similarly to ZINB-WaVE and scVI. This allows for the correction of batch effects if batch annotations are available.
- Remove the sparsity-inducing prior on the loadings matrix. Instead, we introduce sparse loadings via a sparse Gamma prior on them.

Specifically, the model is defined via the following generative process, where we consider gene expression measurements to have been obtained from *B* different experimental batches:

1. For *p* = 1,…, *P* and *j* = 1,…, *K* + *B*, sample a factor loading *w*_*pj*_ ~ Gamma(*β*_1_, *β*_2_).
2. For *n* = 1, …, *N*, sample a scaling factor *l*_*n*_ ~ Gamma (*ν*_1_,*ν*_2_)
3. For *n* = 1,…, *N* and *k* =1,…, *K*, sample a latent factor *z*_*nk*_ ~ Gamma (*α*_1_, *α*_2_).
4. For *n* =1,…, *N* and *p* =1,…, *P*:

a. Sample a raw count *y*_*np*_ ~ Poisson (*l*_*n*_ Σ_*j*_ *w*_*pj*_ [**z**_*n*_, **s**_*n*_]_*j*_).
b. Sample non-dropout event *d*_*np*_ ~ Bernoulli 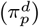.
c. If *d*_*np*_ = 0, *x*_*np*_ = 0. Else, *x*_*np*_ = *y*_*np*_.

The latent variables are **z**, **w**, **d** and **l**. The **z** represents the cells in a lower-dimensional space of size *K* < *P*, **w** is the map from **z** to the observation space, **d** models the occurrence of dropout in each observation and **l** is a cell-specific scaling factor which accounts for capture efficiency and sequencing depth variations.

The batch annotations are included in the model by concatenating them as one-hot encoded vectors **s**_*n*_ (a *B*-dimensional vector where entry *s*_*nb*_ = 1 if cell *n* comes from batch *b*, and is zero otherwise) to the latent space representations **z**_*n*_ and multiplying the resulting vector by the mapping matrix 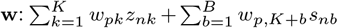. We represent this product as 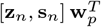. Introducing this additional flexibility in the model results in representations which are disentangled from the experimental batch, thus correcting for batch effects.

Regarding the parameters of the prior distributions - i.e., the model hyperparameters – we choose them so as to warp the space of solutions to the inference problem in a way that favours the structure we believe to describe the data. Namely, in general terms, we aim at obtaining lower-dimensional representations **z**_*n*_ in which similar cell types are clustered together and different ones are well separated. The Normal distribution is usually the distribution of choice for tasks like these, but because we need the Gamma distribution to ensure positiveness, we choose *α*_1_, *α*_2_ to obtain a somewhat similar shape as the normal distribution. In our experiments, we fix *α* = 16 and *α* = 4.

Regarding the factor loadings, we set the shape parameter to be less than 1. In this case, the mass of Gamma distributions concentrates near zero; Ranganath *et al.* (2015) call this type of distribution sparse Gamma and note that it is akin to a soft spike-and-slab prior, which is often used for unsupervised feature selection. This is useful for m-pCMF because it encodes the fact that the latent representations, which should encode relevant biological variability, are only related to a small number of genes. In practice, we fix *β*_1_ = 0.1 and *β*_2_ = 0.3. The prior probabilities 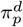 are chosen as in Durif *et al.* (2018), where each 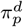 is set to the proportion of non-zeros present in gene *p* across all cells.

Finally, the prior over the scaling factors has a significant impact on the interpretation of m-pCMF. We consider two situations. First, similarly to scVI, we set the scalings to explicitly account for capture efficiency and sequencing depth variations, by making *l*_*n*_ follow the cells’ library sizes. This is done by choosing *ν*_1_ and *ν*_2_ such that the mean and variance of *l*_*n*_ corresponds to the mean *l*_*μ*_ and variance 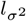 of the library sizes of all cells, respectively. Specifically, this amounts to setting

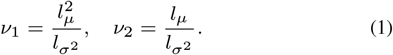

The second situation leverages two (correlated) ideas:

- scRNA-seq data is well described by a zero-inflated negative binomial distribution due to overdispersion;
- The overdispersed counts are largely due to count depth variation.

Specifically, the marginal likelihood of m-pCMF given **z**, **w** and **d** (integrating **l** out) is a zero-inflated negative binomial if each *l*_*n*_ is given by a Gamma distribution with equal prior shape and rate. Formally (and considering only the NB term and one experimental batch for simplicity), this means that if

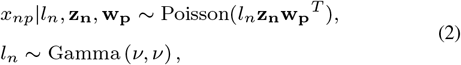

then *x*_*np*_|*l*_*n*_, **z**_**n**_, **w**_**p**_ is NB with mean **z**_**n**_**w**_**p**_^*T*^ and dispersion *ν*. In this construction, *l*_*n*_ has mean of 1 and variance given by 1 /*ν*, and lives in the positive reals. It can be interpreted as a cell size factor. Under this interpretation, we expect variations in *l*_*n*_ to correlate with variations in library sizes. We encode this expectation in the hyperparameter *ν* by setting 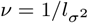, which means that 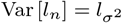.

We have thus defined two ways of encoding library size variations into the model. They both scale the expected gene expression by some expected size factor but, unlike the first one, the second construction ensures that the distribution *p*(*x*_*np*_|**z**_*n*_, **w**_*p*_, *d*_*np*_) is ZINB.

Computing the exact posterior of m-pCMF is computationally intractable, so we approximate it using variational inference (Jordan *et al.*, 1999). In *variational inference* (VI) we posit a family of approximating distributions with known parametric form and optimize its parameters so as to minimize the Kullback-Leibler divergence between the approximation and the true posterior. This minimization is equivalent to maximizing a lower bound on the marginal distribution of the model, i. e., the *evidence lower bound* (ELBO).

Because the cell-specific scalings are Gamma-distributed, the model is, like pCMF, conditionally conjugate and we can derive a *coordinate ascent variational inference* (CAVI) algorithm to update the variational parameters. The batch annotations do not introduce new variables, instead they just increase the dimensions of the loadings matrix. The CAVI algorithm for m-pCMF is thus an extension of the one derived in Durif *et al.* (2018), where we also make use of auxiliary variables *u_npj_* ~ Poisson(*l*_*n*_*c*_*nj*_*w*_*pj*_). The variational approximation is mean-field and the corresponding factors are

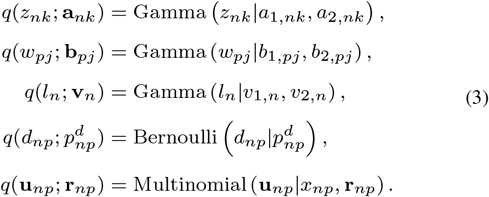

The variational parameters are updated as

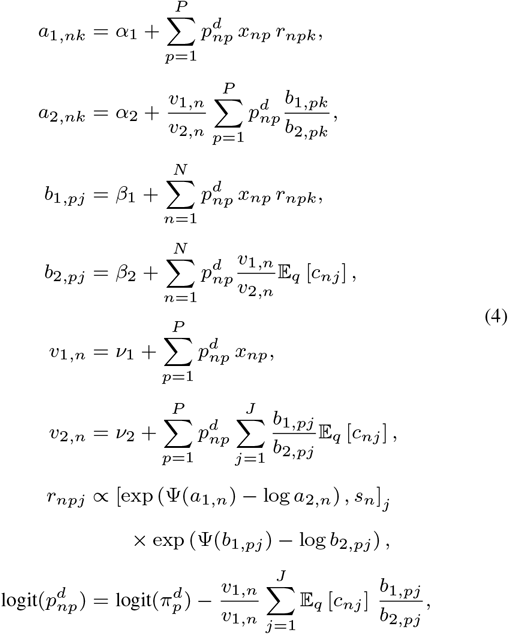

where 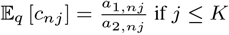 if *j* ≤ *K* and 𝔼_*q*_ [*c*_*nj*_] = *s*_*n,j−*K**_ if *j* > *K*.

Additionally, in order to scale m-pCMF up to large data sets, we turn the CAVI algorithm into an SVI algorithm, following the general procedure described in Hoffman *et al.* (2013). At each iteration we: (1) sample a minibatch of *M* samples from the data set, (2) update the local parameters for each of the samples in the minibatch, and (3) update the global variational parameters **b** using only the local variational parameters for the sampled minibatch. The update of the global parameters starts by computing intermediate global parameters as if each **x**_*m*_ was observed *N* times, and scaling by the mini-batch size *M*. Then, we update **b** at iteration *t* by taking a gradient step.

### 2.2 ZINBayes

We develop a more complex model using the fact that, if

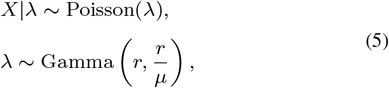

then *X* is NB with mean *μ* and dispersion *r*. We leverage this construction to account for the fact that scRNA-seq data is well described by a ZINB distribution. This model is similar to scVI but it dismisses the use of neural networks and instead performs Bayesian inference over linear mappings from the lower-dimensional space to the count space. This allows for the mappings to be sparse, which can help interpreting what genes are more relevant to what latent component, and to what cell type, consequently.

The following generative process defines the model:

1. For *p* = 1, …, *P*, sample a dispersion parameter *r*_*p*_ ~ Gamma (*θ*_1_, *θ*_2_)
2. For *p* = 1,…, *P* and *j* = 1,…, *K* + *B*:

a. Sample a negative-binomial factor loading *w*_0,*pj*_ ~ Gamma (*β*_1_, *β*_2_).
b. Sample a dropout factor loading *w*_1,*pj*_ ~ Normal (0, 1).
3. For *n* = 1,…, *N*, sample a scaling factor *l*_*n*_ ~ 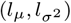
4. For *n* = 1,…, *N* and *k* = 1,…, *K*, sample a latent factor *z*_*nk*_ ~ Gamma (*α*_1_, *α*_1_).
5. Concatenate latent factors **z**_*n*_ with one-hot encoded batch annotation *s*_*n*_: *c*_*nj*_ = [**z**_*n*_, *s*_*n*_]_j_.
6. For *n* =1,…, *N* and *p* =1,…, *P*:

a. Compute a normalized mean gene expression: 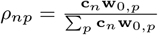
b. Sample a mean gene expression 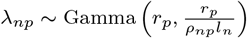
c. Sample a raw count *y*_*np*_ ~ Poisson (λ_*np*_).
d. Sample dropout event *d*_*np*_ ~ Bernoulli (**c**_*n*_**w**_1,*p*_) (logit parameterization).
e. If *d*_*np*_ = 1, *x*_*np*_ = 0. Else, *x*_*np*_ = *y*_*np*_.

There are three main differences between this model and m-pCMF:

1. The negative binomial’s dispersion parameter is independent from the cell-specific scalings, which encode cell size factors. This allows the model to capture other gene-specific dispersion sources besides the count depth variation between cells;
2. The dropout probabilities are related to the latent representations **z**, instead of being entirely separated from the NB part of the generative process;
3. We define the expected relative gene expression frequencies for each cell in *ρ*, which can, in principle, be used for non-biased differential expression. Its scaled version, *ρ***l** gives the NB mean.

We choose the hyperparameters *α*_1,2_ and *β*_1,2_ the same way as described for the corresponding hyperparameters in m-pCMF in Section 2.1, in order to facilitate clustering in **z** and sparse loadings **w**_0_. The prior over the gene-specific dispersion parameters **r** is chosen to be a generic Gamma distribution, as we have no prior expectations of what they should be. However, we do not expect them to be very large (e.g. *r*_*p*_ ≫ 10), so we choose *θ*_1_ = 2, *θ*_2_ = 1.

This model is not conditionally conjugate, due to dropout structure and the parameterization of λ, so we can not derive a CAVI algorithm for the variational updates based on the assumptions described in Hoffman *et al.* (2013). Instead, we use generic methods for inference of all the latent variables. Namely, we use reparameterization gradients-based variational inference to fit the model to data using the Edward probabilistic programming language (Tran *et al.*, 2017).

## 3 Results

In this section we apply the proposed models to real data sets. We evaluate the models in a series of settings: held-out data log-likelihood, dropout imputation, cluster separability in the latent space, separation of biological from technical signals and batch effect correction. As a baseline for a scRNA-seq data-agnostic model, we consider a Factor Analysis (FA).

For FA, we use the scikit-learn Python package (Pedregosa *et al.*, 2011). For ZIFA, we use the Python implementation provided by the authors and the scikit-learn wrapper provided by Lopez *et al.* (2018). For ZINB-WaVE, we use the R package provided by the authors with the additional code provided in Lopez *et al.* (2018) to compute the held-out data log-likelihood. For scVI, we use the Python implementation provided by the authors. For pCMF we use the R package provided by the authors and the scikit-learn wrapper we developed.

In this study we consider data sets with different characteristics in terms of size and protocol used. We are particularly interested in data sets which have been used in previous studies, and whose cells have been previously and reliably annotated. Table 1 summarizes the data sets we consider in this section.

**Table 1.**
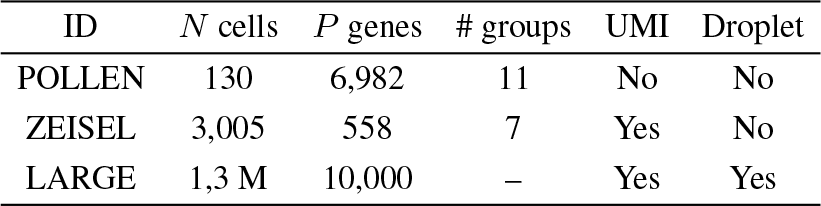
Brief description of the data sets considered in this report. The “UMI” column indicates the use of Unique Molecular Identifiers for the experimental expression quantification. The “Droplet” column indicates whether the data set was generated by a droplet-based experimental protocol. The LARGE data set does not contain annotations.

| ID | $N$ cells | $P$ genes | # groups | UMI | Droplet |
| --- | --- | --- | --- | --- | --- |
| POLLEN | 130 | 6,982 | 11 | No | No |
| ZEISEL | 3,005 | 558 | 7 | Yes | No |
| LARGE | 1,3 M | 10,000 | – | Yes | Yes |

To facilitate comparison with other methods, we utilize the data sets with the same pre-processing used in previous studies. Namely, for POLLEN (Pollen *et al.*, 2014), we use the data available from the *scRNAseq* R package (Risso and Cole, 2016) and retain only the 1000 genes with highest standard deviation as per Risso *et al.* (2018). For ZEISEL (Zeisel *et al.*, 2015), we retain only the 558 genes with highest standard deviation, as in Lopez *et al.* (2018). For LARGE ^1^, we use the count data organized by Lopez *et al.* (2018).

### 3.1 NB improves m-pCMF’s fit

In Section 2.1 we highlighted the fact that setting the hyperparameters of the cell-specific scalings’ Gamma prior as *ν*_1_ = *ν*_2_ yielded a negative-binomial marginal likelihood. In fact, this yields a increase in model fitness, as Fig. 1 illustrates. Specifically, we subsampled 5,000 cells and considered only the 500 most variable genes from the LARGE data set and ran CAVI on m-pCMF five times independently with *ν*_1_ = *ν*_2_ = *ν*, with *ν* equal to the variance of the library sizes, and another five times with *ν*_1_ and *ν*_2_ chosen to achieve a prior mean equal to the mean library size and a prior variance equal to the library size variance, as detailed in Section 2.1.

Because the CAVI algorithm starts with a random initialization of the variational parameters and the ELBO is not convex, each run achieves a different local minimum (this is highlighted in the zoom of the last 300 iterations in the figure). However, the behaviour for the two different m-pCMF settings is consistently different across runs: the NB construction always achieves better values of the objective function than the less expressive Poisson construction. As such, in the subsequent experiments with m-pCMF we always use *ν*_1_ = *ν*_2_ = *ν*, with *ν* equal to the variance of the library sizes.

**Figure 1.**
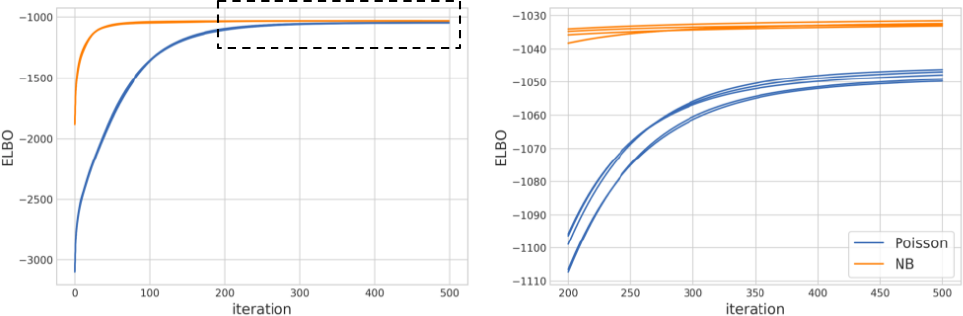
ELBO per iteration of m-pCMF without (Poisson) and with NB structure for 5 runs of 500 iterations. On the left we show the full convergence, and on the right we detail the last 300 iterations.

### 3.2 Effect of the number of Monte Carlo samples for gradient estimation in ZINBayes

Because ZINBayes’ inference is based on (unbiased) estimates of gradients of the objective function, and these estimates are computed via Monte Carlo (MC) sampling, the choice of the number of MC samples has an effect on the variance of these gradient estimates, which in turn impacts the convergence of the inference algorithm. Using the same data as in Section 3.1, we briefly illustrate this impact in Fig. 2.

**Figure 2.**
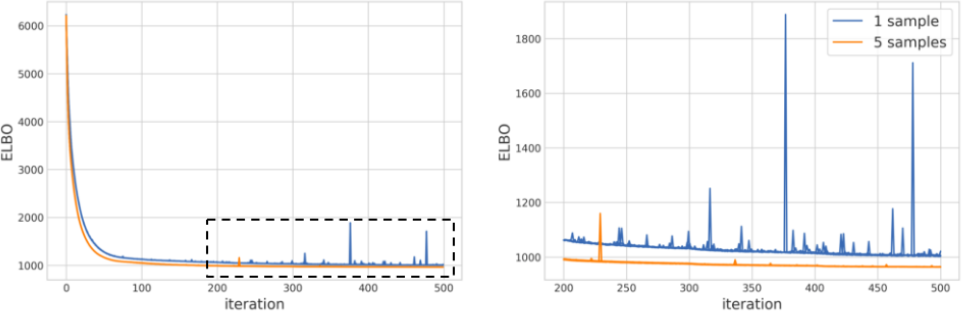
ELBO per iteration of ZINBayes with 1 and 5 Monte Carlo samples for gradient estimation. On the left we show the full convergence, and on the right we detail the last 300 iterations.

In this case, increasing the number of MC samples from 1 to 5 led not only to a better value of the objective function in the fixed number of iterations, but also did so with a considerably lower variance. However, although not shown here, we note that increasing the number of MC samples slows down the algorithm, so there is a trade-off between speed and variance. In practice, we always use 5 MC samples in subsequent experiments.

### 3.3 m-pCMF and ZINBayes disentangle technical factors of variation

Because m-pCMF and ZINBayes account for differences in cell capture efficiency, which lead to differences in library sizes, they can separate these factors of variation from the underlying biological signal. In both models the prior distribution of these scalings are related to the observed library sizes and are thus expected to correlate with them. In Fig. 3 we plot the log of the inferred scalings against the observed log library sizes of the ZEISEL data set. Clearly, the expected correlation is found in these two data sets. Notice that ZINBayes’ scaling factors have the same scale as the observed library sizes, which is expected by their definition. On the other hand, m-pCMF’s scalings, due to the NB construction, vary closer to 1, due to the “exposure” variable interpretation presented in Section 2.1.

**Figure 3.**
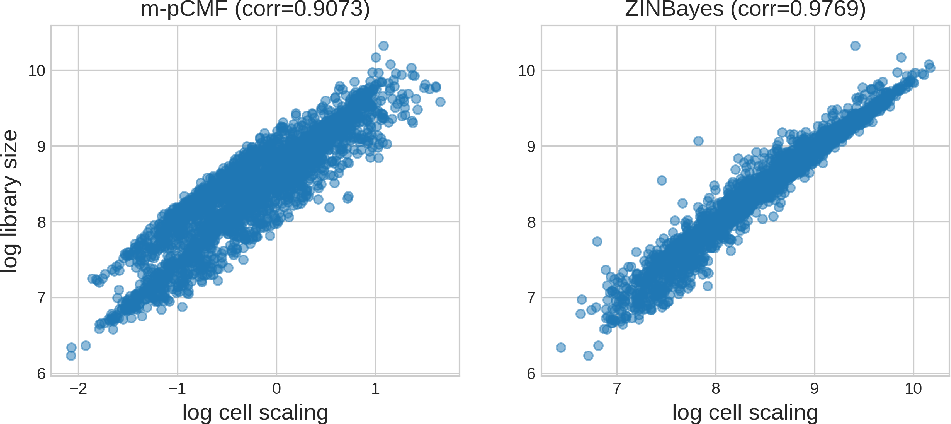
Scatter plots of the log of the estimated cell scaling factors by ZINBayes, m-pCMF and scVI against cell’s log library sizes of the ZEISEL data set. For each scatter plot we compute the corresponding Pearson correlation coefficient. ZINBayes’s scalings correlate very strongly with library sizes.

### 3.4 m-pcmf and ZINBayes are ready for large-scale experiments

Finally, we consider the same LARGE subsample considered in Section 3.1 to note the scalability of m-pCMF and ZINBayes to large data sets. Because these models are ammenable to stochastic variational inference, they can, in principle, be applied to extremely large scRNA-seq data sets with massive numbers of cells. Because these data sets may not fit into memory of the practicioner’s machine, being able to fit an scRNA-seq data model by feeding it mini-batches of data instead of the whole batch of data at once is a valuable feature. While we do not apply our methods to such data sets and always consider, in the subsequent experiments, batch algorithms, we illustrate the capability of m-pCMF and ZINBayes to do so.

Specifically, in Fig. 4 we consider different mini-batch sizes and confirm that, for both models, increasing the mini-batch size leads to higher values of the ELBO, i.e., better model fitness.

**Figure 4.**
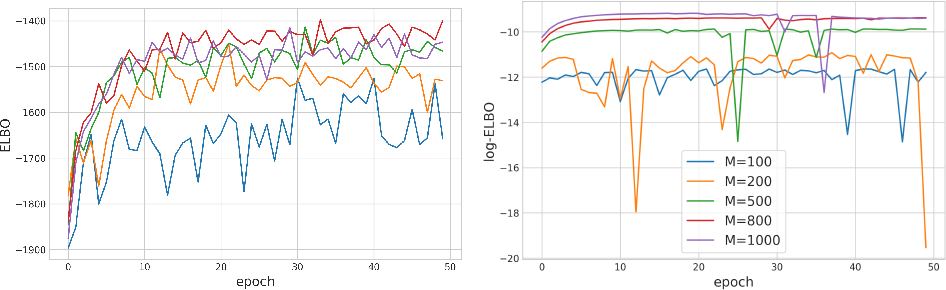
Convergence curves for m-pCMF (left) and ZINBayes (right) with mini-batch stochastic variational inference mechanisms for varying minibatch sizes, M. Each “epoch” consists of a full pass of the data set.

Importantly, we notice that while there is a significant difference in the achieved ELBO values from *M* = 100 to *M* = 500 in the 5,000 cells data set, both for m-pCMF and ZINBayes, the gain decreases significantly as the mini-batch sizes increase from there to *M* = 1000. This means that we may be able to achieve sufficiently good performances at low memory requirements.

### 3.5 Held-out data log-likelihood

We compare the goodness-of-fit and generalization ability of each model by evaluating the per-cell marginal log-likelihood they assign to data unseen during inference. To do this, we perform inference on a large subsample of the data set and then compute the log-likelihood in the rest in a 5-fold cross-validation procedure. We do not consider pCMF as the implementation provided by the authors does not allow for a straightforward way of evaluating the log-likelihood on held-out data.

First, we draw a random subsample of 500 cells from the LARGE data set (containing only the 700 most variable genes) and perform 5-fold cross-validation. The results are shown in Fig. 5. m-pCMF performs worse than all methods except scVI, which yields the worst fit. ZINBayes, while competitive, is unable to beat ZINB-WaVE, ZIFA and FA. Focusing on the test data log-likelihood, scVI is significantly worse than competing methods, despite having been run for a large number of epochs (1000), as recommended by the authors in the case of small data sets. This is related to the fact that there are more local latent variables than global variables in the model, making the use of mini-batch optimization and amortized inference inefficient.

**Figure 5.**
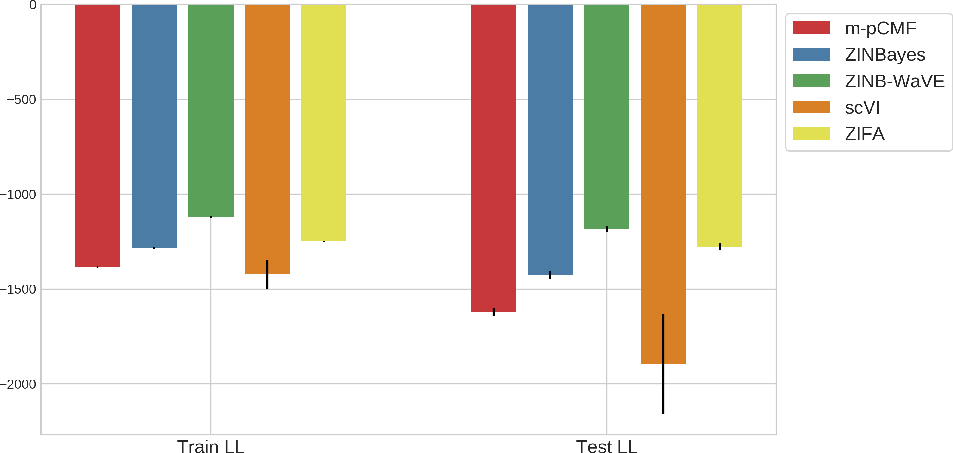
Mean train and test data log-likelihoods of each model with error bars over 5 cross-validation folds in 500 cells from the LARGE data set.

Additionally, we apply the considered scRNA-seq models to different-sized subsets of the LARGE data set and show the log-likelihoods for a held-out test set of 10K cells in Table 2. Here we do not consider a 5-fold CV scheme as ZINB-WaVE and ZIFA become very slow as the number of cells increases. Nevertheless, the obtained values are illustrative of the behaviour of the models with varying number of cells. Namely, scVI has the worst performance for 1K cells and the best performance among all methods for 15K cells. m-pCMF’s performance appears to decrease with the number of cells and ZINBayes’ tendency is to improve.

**Table 2.** Log-likelihood attributed by each model to a held-out set of 10K cells after performing inference on different random subsets of different sizes of the LARGE data set, with 720 genes.

|  | 1K | 5K | 10K | 15K |
| --- | --- | --- | --- | --- |
| m-pCMF | -1098.78 | -1224.10 | -1205.44 | -1462.68 |
| ZINBayes | -1291.83 | -1268.26 | -1251.48 | -1259.26 |
| ZINB-WaVE | -1190.98 | -1173.66 | -1178.52 | -1177.71 |
| scVI | -1509.90 | -1302.81 | -1186.75 | -1193.54 |
| ZIFA | -1276.22 | -1260.26 | -1253.98 | -1272.46 |
| FA | -1211.61 | -1199.72 | -1193.32 | -1196.03 |

### 3.6 Imputation

As noted in Lopez *et al.* (2018), models which attribute a large likelihood to data sets which are dominated by zeros are not necessarily useful for our purpose, as we assume that some percentage of the observed zeros are due to nuisance factors of variation – this was not the case for the data considered in the previous section. As such, to assess model fitness, the authors generated a corrupted training sets by setting 10% uniformly chosen non-zero entries to zero. Then, they fit the perturbed dataset with each of the benchmark methods and evaluate them by comparing the inferred mean values to the original ones, via the median absolute error.

We apply this evaluation procedure to the ZEISEL and POLLEN data sets and show the results in Fig. 6.

**Figure 6.**
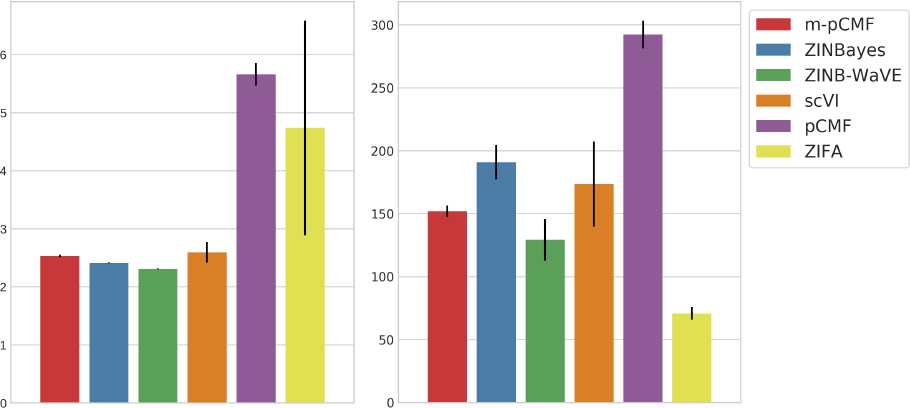
Median imputation error of each model for 5 corruptions of the the ZEISEL (left) and POLLEN (right) data sets.

The results show that pCMF is always the worst performing method. m-pCMF achieves a lower error than scVI on the POLLEN data set, thus corroborating that scVI does not fare well on small data sets. ZINBayes is competitive for the ZEISEL data set.

### 3.7 Latent space structure

Finally, we compare the quality of the structure inferred by each model in the corresponding lower-dimensional space. Similarly to Lopez *et al.* (2018), we consider three metrics for assessing the clustering of cells in the latent space according to the existing biological annotations: Average Silhouette Width (ASW) (Rousseeuw, 1987), Adjusted Rand Index (ARI) (Hubert and Arabie, 1985) and Normalized Mutual Information (NMI) (Vinh *et al.*, 2010). For the ARI and NMI metrics, a clustering is required. In this case, we perform K-means clustering on the latent space with K equal to the true number of clusters. The results for a 5-fold cross-validation for each method in ZEISEL and POLLEN are shown in Fig. 7. For the POLLEN data set, we use the batch annotations to assess whether each model is capable of providing cell-type separability which is independent of the experimental batch of each cell. We evaluate the ASW metric with respect to the batch annotations - in this case, lower is better.

Finally, we consider the POLLEN data set to assess the correlation of the learned lower-dimensional projections with technical factors. To this end, we infer 2-dimensional projections of the data and evaluate the correlation of each dimension of each model’s latent space with the library size and detection rates of the cells. The results are shown in Fig. 8. The ASW of ZINBayes’s latent space does not have a signficant change from 10D (Fig. 7, right panel) to 2D. Furthermore, ZINBayes is the method that achieves better clustering of different cell types while maintaining a very low correlation with library sizes and a median correlation with detection rates. Notably, FA, ZIFA and pCMF’s latent factors are not only unable to cluster cell types, but the latent structure is also extremely correlated with library sizes.

### 3.8 Discussion

In terms of model fitness, the results show that ZINBayes outperforms scVI for the 500, 1K and 5K cell-subsamples of the LARGE data set, the ZEISEL data set (both in terms of marginal log-likelihood and imputation). m-pCMF outperforms scVI in the 500, 1K and 5k cell-subsamples of the LARGE data set and the POLLEN data set (imputation). All models yield a better fit than pCMF. ZINB-WaVE is always the best-fitting model, except for the POLLEN data, for which ZIFA yields the lowest imputation error.

The fact that ZINB-WaVE is based on a similar parameterization to scVI withouth the non-linearities but still provides a better overall fit suggests that either the non-linearities are not necessary, its fitting mechanism is able to achieve better solutions of the parameters, or scVI’s prior hyperparameters are inadequate. In this sense, the fact that ZINBayes also dismisses the use non-linearities points towards either the second or third hypothesis. To further investigate this, we should analyse the estimated values of ZINB-WaVE’s parameters to see if they can not be captured by the prior distributions in ZINBayes.

In terms of cell type separability in the reduced spaces, m-pCMF and ZINBayes yield higher ASW scores and moderately lower ARI and NMI scores than all competing methods, except pCMF, in the ZEISEL data set. For the POLLEN data set, ZINBayes provided a better biological structure than all the other models according to every metric. This suggests that ZINBayes is a better suited model for smaller data sets than competing methods. Typically, smaller data sets exhibit a much larger gene-specific dispersion than droplet-based cohorts, and ZINBayes’ prior on the over-dispersion is generic enough to allow for a wide range of values for this parameter while still constraining the parameter space, which facilitates optimization. This characteristic may also explain why m-pCMF performs very poorly on the POLLEN data set, being unable to find any biological structure, as it is unable to model gene-specific dispersion.

In terms of correction of batch effects, we observed for the POLLEN data set that only ZINBayes’ and ZINB-WaVE’s latent structures were uncounfounded by the cells’ experimental batches. While for ZINBayes we explicitly included the batch annotations in the model, ZINB-WaVE was able to achieve this good performance without any batch information. We showed that these were the two models whose latent components were less correlated with library size variation, which was the main counfounding factor between experimental batches in the POLLEN data set.

## 4 Conclusion

Two novel generative models for dimensionality reduction of scRNA-seq data were proposed. ZINBayes shares some structure with current state-of-the-art models, which may explain the overall better performance compared to m-pCMF, which, for example, does not consider gene-specific over-dispersion. While none of the proposed models provides an overall improvement in performance over the state-of-the-art methods ZINB-WaVE and scVI, we note that: 1) ZINBayes’ performance on data sets with for which the number of cells is not large compared to the number of genes does not decrease, contrary to scVI; and 2) m-pCMF and ZINBayes do not need the full batch of data to perform inference, contrary to ZINB-WaVE.

Future work would first consist of performing more extensive experiments on m-pCMF and ZINBayes. For example, we did not analyse the factor loadings on which sparsity was imposed through the sparse Gamma prior. These loadings may provide information on the contribution of genes to the lower-dimensional representations and could be related with cell types, as done in Buettner *et al.* (2017). Furthermore we did not assess the posterior uncertainties which could be useful, for example, in differential expression analysis.

We argue that these scalable generative models will be useful in unlocking the many potential applications of scRNA-seq by providing a detailed look into the cell-to-cell heterogeneities in terms of gene expression. Characterizing cells in these terms can be used in describing diseases in terms of, for example, the presence of a certain kind of cells which is a step forward towards personalized medicine.

**Figure 7.**
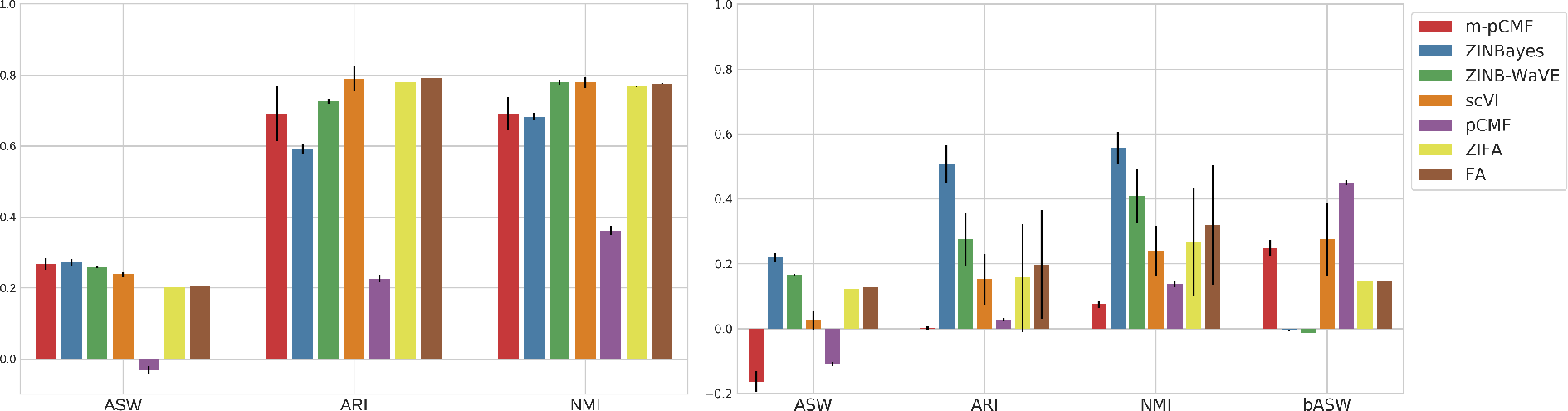
Clustering metrics on the latent space for 5 runs of each model on the whole ZEISEL (left) and POLLEN (right) data sets. bASW is the ASW score with respect to batch annotations.

**Figure 8.**
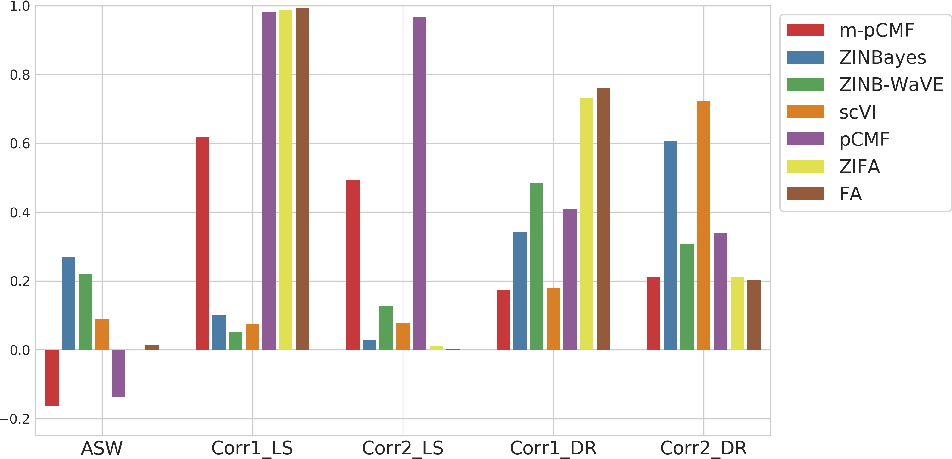
ASW of the 2-dimensional latent representations with respect to cell types of the POLLEN data set and absolute Pearson correlation coefficients of each dimension with library sizes (Corr1_LS, Corr2_LS) and detection rates (Corr1_DR, Corr2_DR).

## Acknowledgements

This work was partially supported by the European Union Horizon 2020 research and innovation program under grant agreement No. 633974 (SOUND project), and also by Fundação para a Ciência e a Tecnologia - FCT under contracts INESC-ID (UID/CEC/50021/2013), IT (UID/EEA/50008/2013), IDMEC/LAETA (UID/EMS/50022/2013), and projects PERSEIDS (PTDC/EMS-SIS/0642/2014) and PREDICT (PTDC/CCI-CIF/29877/2017).

## Footnotes

1 10X Genomics, https://support.10xgenomics.com/single-cell-gene-expression. Accessed 2018-06-10.

